# Directed conversion of porcine extended pluripotent stem cells into trophoblast-like stem cells through modulation of conserved TGF-β and ERK signaling pathways

**DOI:** 10.64898/2026.02.04.703856

**Authors:** Chi-Hun Park, Young-Hee Jeoung, JiTao Wang, Bhanu P. Telugu

**Author notes:** To whom correspondence should be addressed: Bhanu Telugu, Division of Animal Sciences, University of Missouri, Columbia, MO 65211.

## Abstract

Trophoblast stem cells (TSCs) provide a tractable system for interrogating the signaling pathways that govern extraembryonic lineage commitment. Although trophoblast specification has been extensively characterized in humans and rodents, comparable tools and molecular frameworks remain poorly defined in pigs. Here, we identify defined biochemical conditions that enable the conversion of porcine extended pluripotent stem cells (pEPSCs) into TSCs. Pharmacological inhibition experiments demonstrate that coordinated repression of TGF-β/Activin and MEK/ERK signaling is sufficient to induce and maintain a stable trophoblast transcriptional program. Under these conditions, cells robustly upregulate core trophoblast regulators, including *CDX2, GATA3, KRT7/18, HAND1*, and *ELF5*, while concomitantly suppressing pluripotency- and hypoblast-associated gene networks. Bulk transcriptomic profiling reveals extensive lineage reprogramming, with enrichment of pathways related to cell adhesion, extracellular matrix organization, and placental development. Functional *in vivo* assays further show that induced trophoblast-like cells form small, non-teratomatous lesions that express extraembryonic markers, whereas parental pEPSCs generate teratomas that contain derivatives of all three germ layers. Together, these findings establish that combined inhibition of TGF-β/Activin and MEK/ERK signaling is sufficient to specify porcine trophoblast identity from pluripotent stem cells and provide a biochemical framework for dissecting conserved and species-specific mechanisms underlying trophoblast specification and placental development.

**Highlights:**

- Defined signaling conditions enabling stable conversion of porcine extended pluripotent stem cells (EPSCs) into trophoblast-like stem cells (TSC).
- Dual inhibition of TGF-β/Activin and ERK pathways drives robust trophoblast commitment.
- Transcriptional reprogramming reveals conserved trophectoderm regulatory networks distinct from pluripotency and hypoblast states.
- Induced porcine TSCs display restricted *in vivo* potential, consistent with trophoblast identity.

## Introduction

The placenta, a defining feature of eutherian mammals, is essential for fetal growth due to its roles in nutrient exchange, endocrine signaling, and immune regulation (1-3). While these core functions of the placenta are conserved, the cyto-architecture and the degree of maternal-fetal interaction differ substantially among species (4). A particularly notable distinction is the extent of trophoblast invasion into maternal tissues. In humans and rodents, a hemochorial placenta forms, characterized by trophoblast cell invasion of the maternal endometrium and direct contact with maternal blood (5). Conversely, pigs develop a non-invasive epitheliochorial placenta that preserves the maternal epithelium, stroma, and vasculature throughout gestation (6, 7). These anatomical differences highlight the necessity for species-specific models to advance the understanding of placental development and function.

In the human placenta, trophoblasts differentiate into several specialized subtypes that collectively establish and maintain the maternal-fetal interface. Cytotrophoblasts (CTBs) represent a proliferative, stem-like population that can fuse to form multinucleated syncytiotrophoblasts (STBs), which are responsible for hormone production and nutrient exchange. A subset of CTBs undergoes epithelial-to-mesenchymal transition to generate extravillous trophoblasts (EVTs), which invade maternal tissues and remodel uterine spiral arteries to facilitate placental perfusion (8-10). In contrast, the diversity and lineage organization of trophoblast cells in pigs are less well characterized. Molecular profiling has revealed region-specific expression of transporters and signaling molecules within the porcine trophoblast, indicating functional heterogeneity and the existence of specialized subtypes. Notably, areolar trophoblast cells at uterine gland openings display unique molecular signatures and are involved in the uptake of glandular secretions (11, 12). However, the absence of robust experimental systems has hindered mechanistic studies of porcine trophoblast development and function. Although trophoblast stem cell (TSC) lines have been derived from pre-/peri-implantation pig embryos (<u>Edison</u>) (13, 14), their application has been limited by issues of limited characterization, rigor, and reproducibility.

During early mammalian development, the first lineage-specification decision results in the separation of blastomeres into trophectoderm (TE) and inner cell mass (ICM), with the ICM subsequently differentiating into epiblast and hypoblast (primitive endoderm). In several species, defined culture conditions enable the derivation of TSCs from blastocysts, stem cells or placental tissues (15). Studies in humans have shown that both naïve and primed pluripotent stem cells (PSCs) can be directly converted into TSCs under specific signaling conditions (15-17). In pigs, recent progress has led to the derivation of pluripotent stem cells (pPSCs) (18-20), specifically the extended pluripotent stem cells (pEPSCs) which have demonstrated potential for trophoblast-like differentiation (21, 22). Despite these advances, the lack of robust and reproducible *in vitro* models continues to hinder systematic investigation of porcine trophoblast lineage specification and maintenance. This study hypothesizes that pPSCs can be directly converted into authentic TSCs with sustained self-renewal and stable lineage identity, with the ultimate goal of uncovering conserved signaling pathways required for efficient conversion.

### Induction of trophoblast differentiation from porcine EPSCs

Porcine EPSC (pEPSC) were derived from day 7 *in vivo* blastocysts using the previously reported 3iLAF condition(18) (Supplementary Fig. 1A). The established cell lines exhibited a compact, undifferentiated morphology, strong alkaline phosphatase activity (Fig. 1A), and expression of core pluripotency markers such as OCT4, SOX2, NANOG, and SSEA4, as confirmed by immunofluorescence (Fig. 1B; Supplementary Fig. 1B). The capacity of pEPSCs for trophoblast conversion was evaluated using two media previously shown to induce trophoblast identity in human pluripotent stem cells (PSCs): BAP medium (BMP4, A83-01, PD0325901 or PD173074) (17,18) and CTB medium (A83-01, CHIR99021, EGF, and ascorbic acid) (15). BAP medium did not support trophoblast induction, resulting in minimal epithelial transition and extensive cell death within 48 hours. The FGFR inhibitor PD173074 was cytotoxic even at low concentrations, leading to rapid loss of viable cells. In contrast, CTB medium induced a rapid morphological transition, with colonies flattening and adopting a trophoblast-like epithelial morphology within three days (Fig. 1C). The composition of the basal medium was a critical determinant of cell survival and conversion efficiency. Defined, N2B27-based formulations were ineffective, while substitution with DMEM/F12 supplemented with 5% knockout serum replacement (KSR) or fetal bovine serum (FBS) significantly improved cell survival and epithelial organization under BAP conditions (Supplementary Fig. 1C). These results indicate that both growth factor signaling and basal nutrient composition strongly influence trophoblast induction from pEPSCs. After serial passaging, stable epithelial cultures were established that showed robust upregulation of the pan-trophoblast marker CDX2 and the mononuclear trophoblast regulator GATA3, along with repression of pluripotency genes (OCT4, NANOG) and hypoblast-associated markers (GATA6, SOX17) (Fig. 1D,1E). Collectively, these results demonstrate successful commitment of pEPSCs to the trophoblast lineage under defined induction conditions.

**Figure 1.**
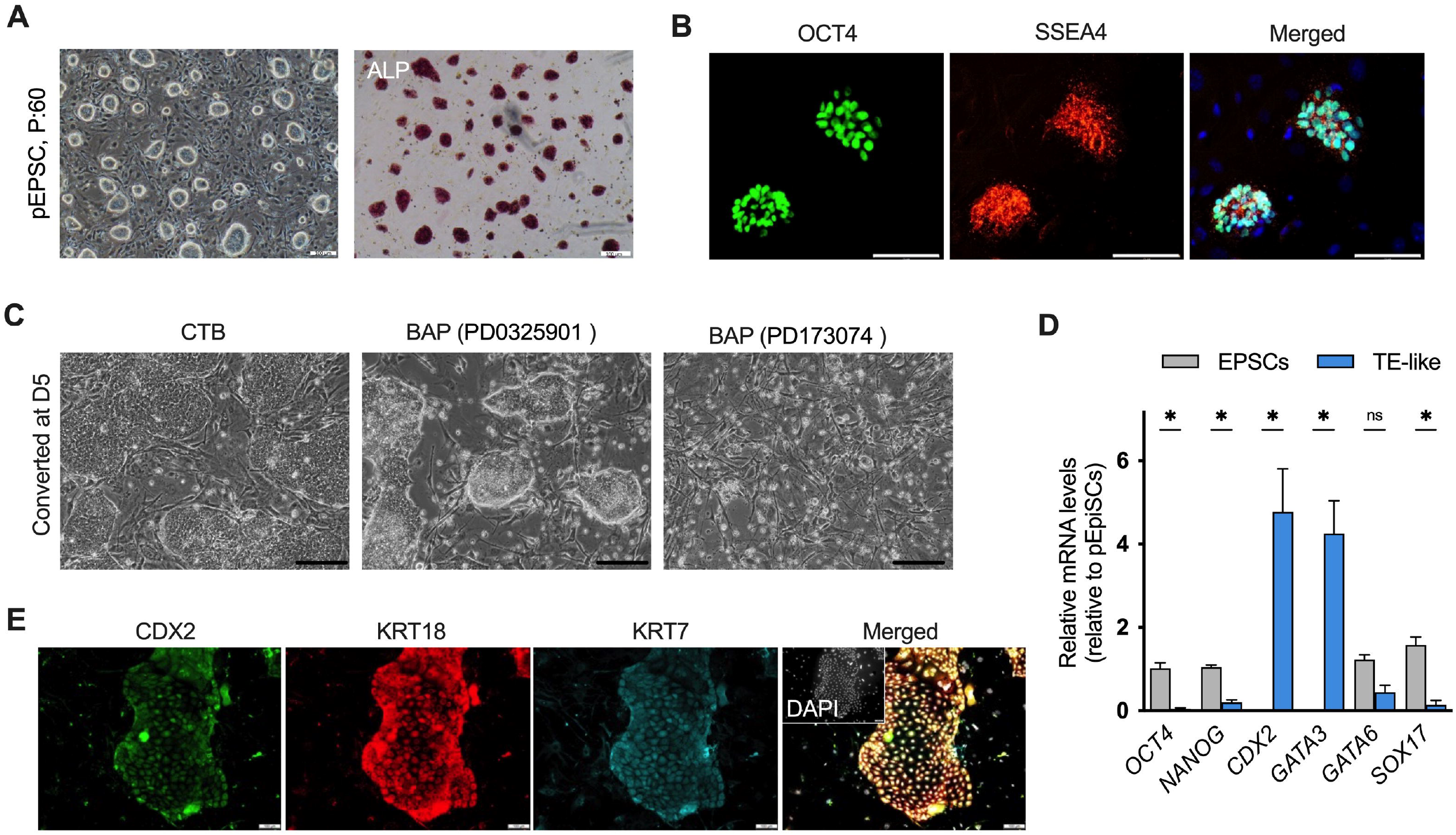
Derivation and characterization of porcine epiblast stem cells (EPSCs) and their conversion toward a trophoblast stem cell (TSC)-like state. (A) Representative phase-contrast image of porcine EPSC colonies at passage 60 displaying typical epithelial morphology (right) and alkaline phosphatase (ALP) staining confirming pluripotent identity (left). Scale bars, 100 μm. (B) Immunofluorescence staining of EPSCs showing robust nuclear expression of the pluripotency markers OCT4 and SSEA4. Merged images include DAPI nuclear counterstaining. Scale bars, 75 μm. (C) Morphological changes during early trophoblast induction under human TSC culture conditions. At day 5, EPSCs maintained in control conditions (CT) retain epiblast-like morphology, whereas combined inhibition of TGF-β/Activin and MEK/ERK signaling (BAP) using PD0325901 or PD173074 induces epithelial colonies with trophoblast-like features. Scale bars, 100 μm. (D) Quantitative RT–PCR analysis of lineage marker expression in EPSCs (gray) and induced TE-like cells (blue), normalized to EPSC controls. Pluripotency-associated genes (*OCT4, NANOG*) are reduced, while trophoblast-associated markers (*CDX2, GATA3*) are significantly upregulated. *GATA6* shows no significant change, and *SOX17* remains low. Data are presented as mean ± SEM; *P < 0.05; ns, not significant. (E) Immunofluorescence analysis of induced TE-like colonies showing expression of trophoblast markers CDX2, KRT18, and KRT7. Merged images include DAPI nuclear staining. Scale bars, 100 μm.

### Signaling requirements for trophoblast induction

To identify the signaling pathways involved in trophoblast induction, pEPSCs were subjected to defined perturbations targeting the TGF-β/Activin and MEK/ERK pathways and analyzed during early stages of derivation (P0-P3). Inhibition of TGF-β/Activin signaling with either A83-01 or SB431542 efficiently suppressed pluripotency-associated transcripts (*OCT4, SOX2, NANOG*) and hypoblast-related genes (*SOX17, GATA6*) (Supplementary Fig. 2A), and induced key trophoblast regulators such as *CDX2, GATA3, KRT8*, and *KRT18* (Fig. 2A). These findings indicate that TGF-β/Activin inhibition is essential for initiating trophoblast reprogramming by day 5 (Supplementary Fig. 2B). Combined inhibition of TGF-β/Activin and MEK/ERK signaling further enhanced both induction efficiency and stabilization of trophoblast-specific transcriptional programs. Among the tested conditions, dual treatment with A83-01 and PD0325901 (A/P) produced the most robust and coordinated activation of the trophoblast gene network. At P0, this condition induced the highest levels of *CDX2*, together with downstream trophoblast regulators such as *ELF5* and *GATA3*, consistent with synergistic effects on early trophoblast specification.

**Figure 2.**
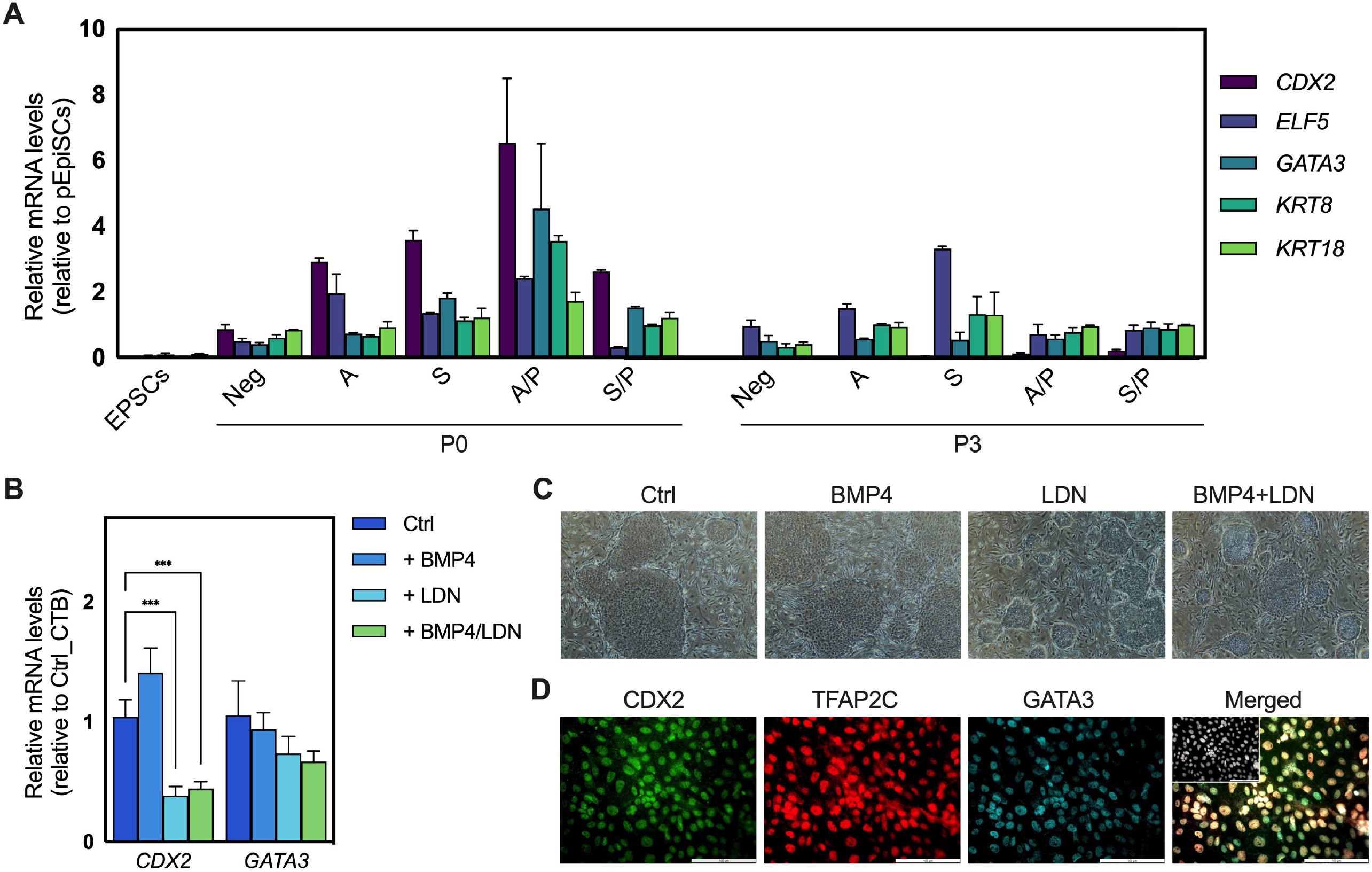
Signaling requirements for trophoblast induction from porcine EPSCs. (A) Quantitative RT–PCR analysis of trophoblast-associated gene expression during early derivation (P0, day 5) and after stabilization (P3) under defined inhibitor conditions. Porcine EPSCs were cultured with TGF-β/Activin pathway inhibitors (A83-01 or SB431542), MEK/ERK inhibition (PD0325901), or their combinations. Expression levels of CDX2, ELF5, GATA3, KRT8, and KRT18 are shown relative to parental EPSCs. (B) Quantitative RT–PCR analysis of CDX2 and GATA3 expression in induced TSC-like cells cultured with BMP4, the BMP inhibitor LDN193189 (LDN), or their combination, relative to control TSC conditions. BMP4 treatment significantly increased CDX2 expression, whereas BMP inhibition reduced CDX2 levels. GATA3 expression showed minimal variation across conditions. Data are presented as mean ± SEM; ***P < 0.001. (C) Representative phase-contrast images of induced cultures at day 5 under control conditions, BMP4 supplementation, BMP inhibition (LDN), or combined BMP4 + LDN treatment. Cultures treated with BMP4 exhibited more compact epithelial morphology, whereas LDN-treated groups showed reduced trophoblast-like features. (D) Immunofluorescence staining of induced TSC-like cells showing nuclear expression of trophoblast transcription factors CDX2, TFAP2C, and GATA3, with merged images including DAPI nuclear counterstaining. Scale bars, 100 μm.

The contribution of BMP signaling during induction was evaluated by adding BMP4 or the BMP inhibitor LDN193189 to the control condition containing PD0325901 (23). BMP4 treatment significantly increased *CDX2* expression, while BMP inhibition markedly reduced *CDX2* levels (Fig. 2B). In contrast, *GATA3* expression remained largely unchanged across conditions, indicating that BMP signaling selectively enhances early TE-associated transcription but is not strictly required for trophoblast lineage establishment. Morphologically, BMP4-treated cultures exhibited more compact epithelial organization, whereas LDN-treated cells displayed reduced trophoblast-like features (Fig. 2C). Together, these findings demonstrate that TGF-β/Activin inhibition is necessary for trophoblast reprogramming, while concurrent MEK/ERK inhibition enhances induction efficiency and stabilizes trophoblast identity (Fig. 2D) (15, 24). BMP signaling has been shown to play a supportive, but non-instructive role by improving derivation efficiency.

### Key transcriptional dynamics during early conversion

After establishing optimized induction conditions (CTB medium supplemented with PD0325901 and BMP4), the transcriptional dynamics underlying early trophoblast conversion were examined. Consistent with morphological observations, pEPSC colonies rapidly transitioned from compact, dome-like structures to flattened epithelial sheets during the initial days of induction (Fig. 3A). Time-course qPCR analysis revealed a hierarchical activation of trophoblast markers rather than their simultaneous induction (Fig. 3B). *CDX2* was the earliest transcription factor to be upregulated, preceding the activation of other trophoblast regulators. *GATA3* expression increased by day 3, followed by induction of *ELF5*, consistent with progressive consolidation of trophoblast identity. Structural markers *KRT8* and *KRT18* increased more gradually, reflecting epithelial maturation rather than initial lineage commitment. In contrast, *TEAD4* expression declined transiently during early induction before re-emerging at later stages, suggesting dynamic remodeling of trophoblast regulatory circuits during the transition from pluripotency to a stabilized stem-like state (25). *HAND1* expression was detected predominantly at later stages, consistent with its association with differentiated trophoblast sub-lineages. *ETS2* expression decreased upon induction, consistent with its context-dependent and non-essential role in TSC maintenance (26).

**Figure 3.**
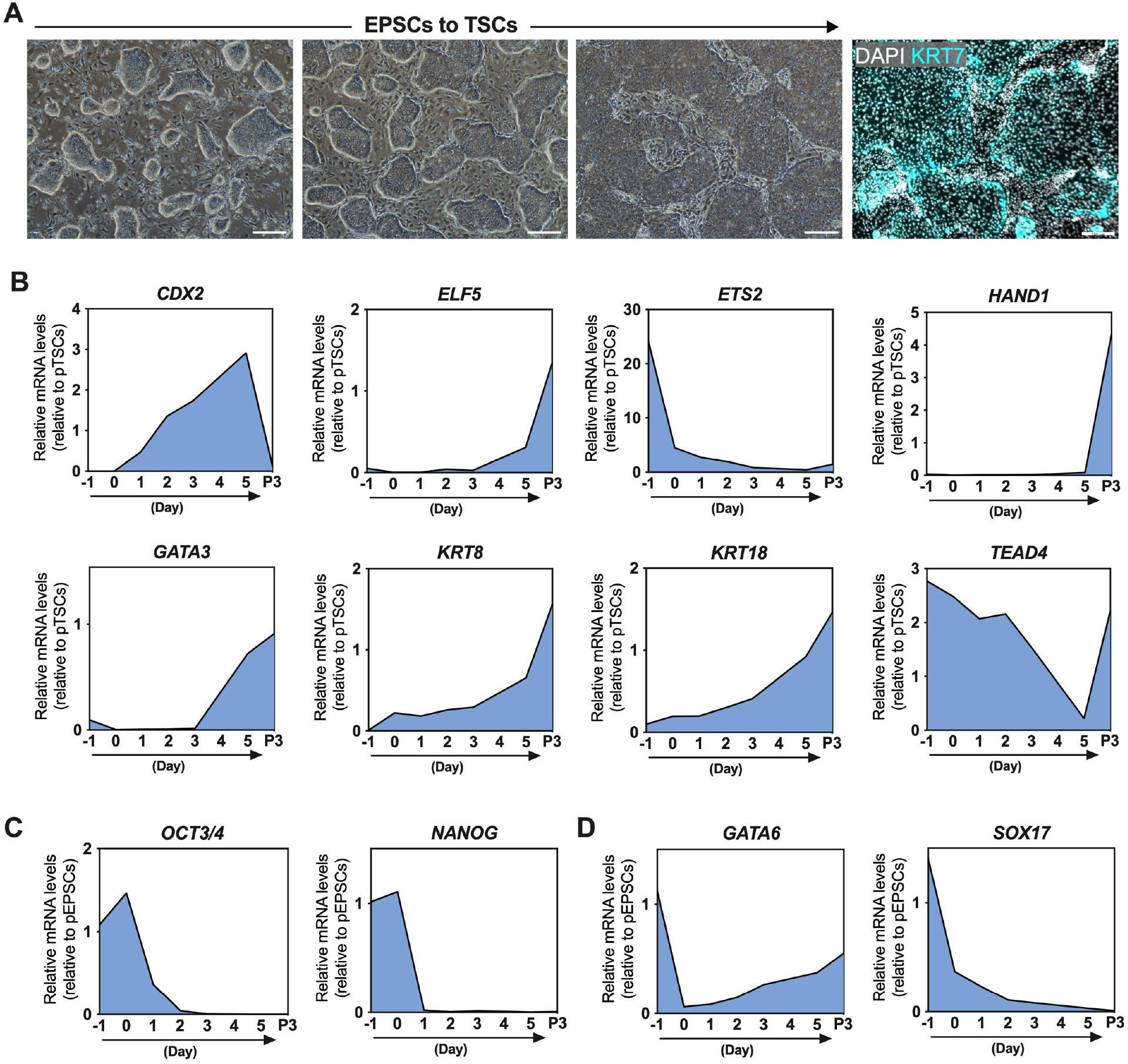
Temporal transcriptional dynamics during conversion. (A) Representative phase-contrast images illustrating morphological progression during EPSC-to-TSC conversion. Immunofluorescence staining at the end of induction confirms expression of the trophoblast marker KRT7 (cyan) with DAPI nuclear counterstaining. (B) Time-course quantitative RT–PCR analysis of trophoblast-associated transcription factors and epithelial markers during induction, including *CDX2, ELF5, ETS2, HAND1, GATA3, KRT8, KRT18*, and *TEAD4*. Expression levels are shown relative to established porcine TSCs (pTSCs). (C) Temporal expression of pluripotency-associated genes *NANOG, OCT3/4*, and *SOX2* during induction, normalized to EPSC controls, demonstrating rapid downregulation of core pluripotency factors. (D) Expression dynamics of lineage-associated markers *EOMES, GATA6*, and *SOX17* during conversion, indicating suppression of mesendodermal and hypoblast-associated programs. (E) Growth kinetics of EPSCs-p:65 (gray) and induced TSC-like cells (iTSCs-p:15; blue) following plating, showing comparable proliferative capacity during early passages. Scale bars, 100 μm.

Pluripotency genes *OCT4, SOX2*, and *NANOG* were rapidly and uniformly downregulated (Fig. 3C), indicating efficient exit from the pluripotent state. Hypoblast-associated genes *SOX17* and *EOMES* were sharply suppressed and remained low throughout induction, while *GATA6* showed an initial decrease followed by partial re-expression at later stages (Fig. 3D; Supplementary Fig. 3A). Immunofluorescence analysis confirmed detectable GATA6 protein in a subset of induced cells (Supplementary Fig. 3B). Notably, GATA6 expression was also observed in allantochorionic extraembryonic tissues of day-25 porcine conceptuses (Supplementary Fig. 3C), suggesting context-dependent expression in porcine extraembryonic lineages. Collectively, these data demonstrate a stepwise regulatory process underlying trophoblast induction, characterized by rapid exit from pluripotency and sequential activation of conserved trophoblast stem cell regulators. The preservation of core factors such as *CDX2, GATA3*, and *ELF5* supports the existence of shared regulatory modules governing TSC identity across mammals.

### Long-term culture and substrate-dependent maintenance of pTSCs

To establish conditions supporting long-term and stable maintenance of pTSC-like cells, feeder-based induction was compared with subsequent feeder-free culture on defined extracellular matrices (Fig. 4A). During early induction, culture on MEF feeders promoted efficient attachment and survival, facilitating the establishment of epithelial colonies. After serial passaging, feeder-free culture on Matrigel supported more uniform epithelial morphology and sustained proliferation. Withdrawal of PD0325901 following induction led to pronounced changes in growth behavior and epithelial organization (Fig. 4B). Cultures maintained without PD0325901 adopted a flattened epithelial morphology resembling human CTBs, while continued PD0325901 treatment preserved a more compact epithelial architecture. Both conditions retained expression of canonical trophoblast markers, including KRT7 and KRT18 (Fig. 4C), indicating that trophoblast identity was maintained independent of ERK inhibition.

**Figure 4.**
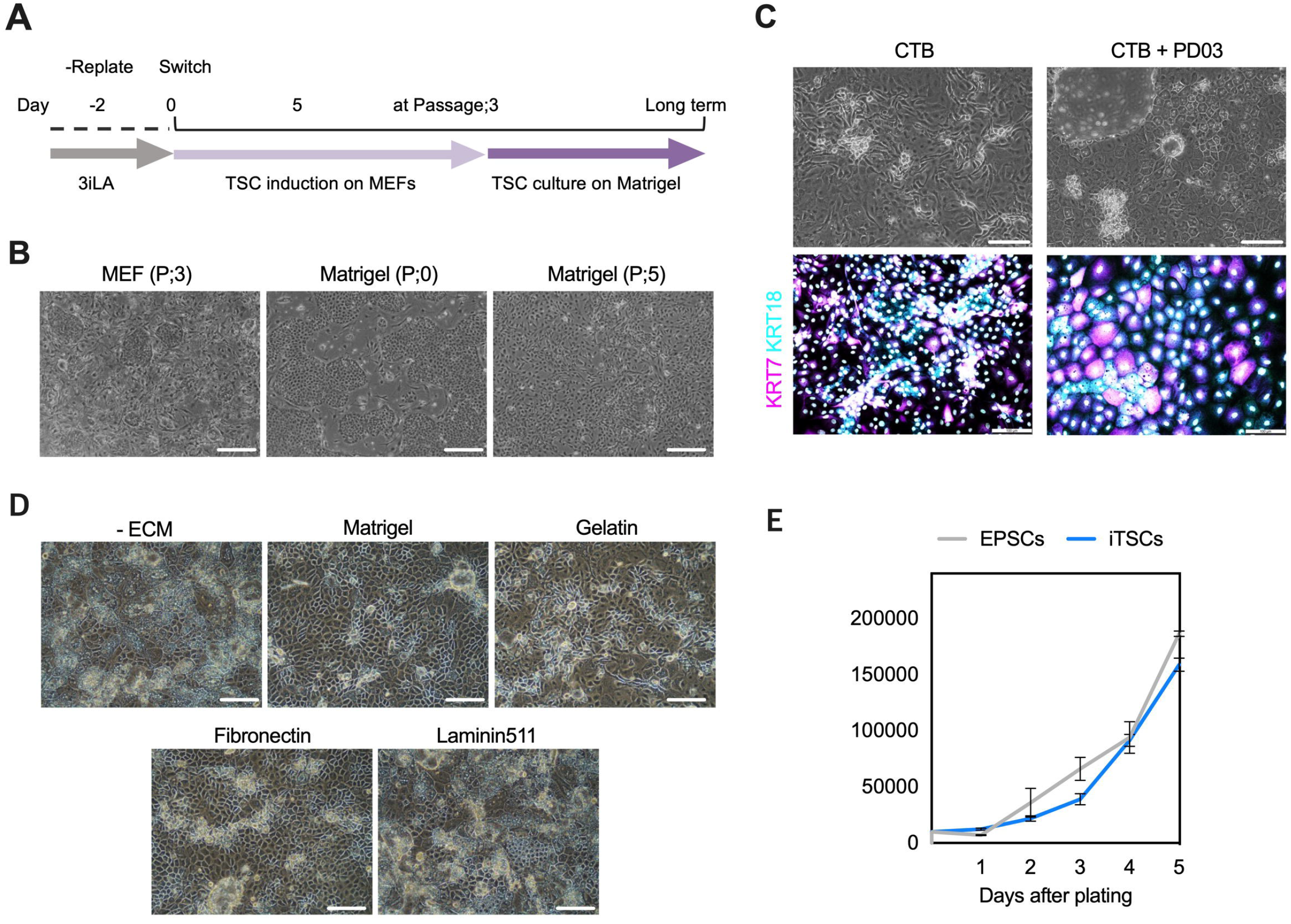
Long-term culture and extracellular matrix requirements for maintenance of porcine TSC-like cells. (A) Schematic overview of the induction. Porcine EPSCs were induced toward the trophoblast lineage on MEF feeders, and subsequently transitioned to feeder-free culture on Matrigel. (B) Representative phase-contrast images showing morphological changes following transfering from the feeder to Matrigel culture (C) Morphological and immunofluorescence characterization of TSC-like cells. Phase-contrast images and immunostaining for KRT7 and KRT18 demonstrate retention of trophoblast, with or without PD0325901. PD0325901 resulted in slower growth and maintained compact epithelial organization. (D) Evaluation of extracellular matrix (ECM) requirements for feeder-free maintenance. Cells were cultured on Matrigel (1%), gelatin (0.1%), fibronectin (1 µg/cm^2^), laminin-511 (1 µg/cm^2^), or without ECM. Matrigel supported the most homogeneous epithelial morphology, whereas other substrates promoted differentiation and heterogeneity. (E) Proliferation kinetics of induced TSC-like cells compared with parental EPSCs over five days following plating, showing comparable growth rates under optimized feeder-free conditions.

Transitioning from feeder-based to feeder-free Matrigel culture led to progressive morphological homogenization across successive passages. Long-term Matrigel-maintained cultures formed cohesive epithelial sheets with reduced heterogeneity, consistent with stabilization of a TSC-like state independent of feeder-derived cues. Among the tested substrates, Matrigel, gelatin, and fibronectin effectively supported sustained growth and epithelial organization, while laminin-511, or matrix-free conditions promoted differentiation and morphological heterogeneity (Fig. 4D). Under optimized feeder-free conditions, pTSC-like cells displayed proliferation kinetics comparable to parental pEPSCs (Fig. 4E), indicating robust self-renewal capacity.

### Transcriptomic validation of lineage conversion

Bulk RNA-seq analysis comparing parental porcine EPSCs and derived induced (i) pTSCs revealed extensive transcriptional reprogramming accompanying lineage conversion. Principal component analysis demonstrated clear segregation of iTSCs from parental EPSCs, with iTSCs clustering distinctly from both pluripotent EPSCs and extraembryonic endoderm (XEN) cells derived from a previously published dataset (27), indicating acquisition of a unique transcriptional identity rather than partial lineage overlap (Fig. 5A). Consistent with this lineage separation, a focused heatmap revealed strong upregulation of key trophoblast-associated genes, such as *CDX2, GATA2/3, ELF5, TFAP2A, KRT8*, and *KRT18* in TSCs relative to EPSCs (Fig. 5B), further supported by differential gene expression analysis showing widespread transcriptomic changes across thousands of genes (Supplementary Fig. 4A). Gene Ontology (GO) and KEGG pathway enrichment analyses highlighted distinct lineage-specific functional signatures. Genes upregulated in parental pEPSCs were enriched for pathways associated with pluripotency and epiblast identity, including transcriptional regulation, developmental signaling, and chromatin organization. Correspondingly, GO terms such as “regulation of transcription, DNA-templated” and “cell fate commitment,” along with enrichment of nuclear and chromatin-associated cellular components and DNA-binding transcription factor activity, were prominent in EpiSCs (Supplementary Fig. 4B,C). In addition, genes upregulated in TSCs exhibited significant enrichment of KEGG pathways related to trophoblast function, including PI3K-Akt signaling, focal adhesion, ECM-receptor interaction, lysosome, and sphingolipid signaling (Fig. 5C). GO biological process categories such as “extracellular matrix organization,” “cell adhesion,” “vesicle-mediated transport,” and “apoptotic cell clearance,” as well as cellular component enrichment in membrane-associated and lysosomal compartments, were selectively enriched in pTSCs (Supplementary Fig. 4C). In addition, pathway-level analysis revealed differential utilization of receptor tyrosine kinase-associated signaling modules between EPSCs and pTSCs (Supplementary Fig. 4D).

**Figure 5.**
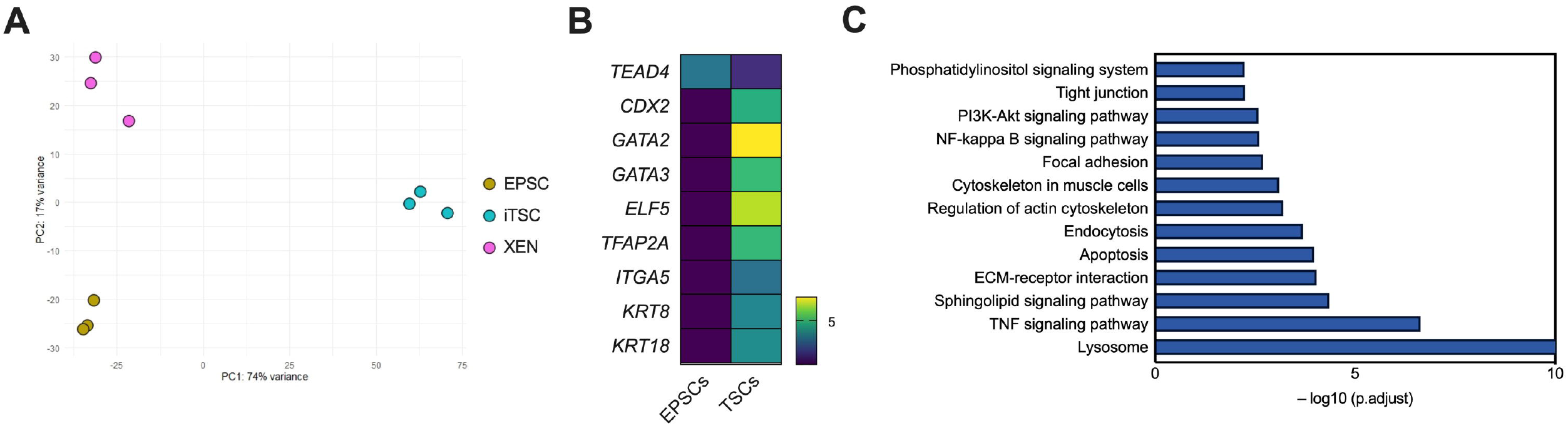
Transcriptomic profiling of porcine EpiSCs and induced TSCs. (A) Principal component analysis (PCA) of global gene expression profiles from parental porcine EpiSCs, induced TSCs (iTSCs), and extraembryonic endoderm (XEN) cells derived from a previously published dataset (GSE128149). iTSCs cluster distinctly from EpiSCs and are clearly separated from XEN cells, indicating acquisition of a transcriptional identity distinct from pluripotent and endodermal lineages. (B) Heatmap showing relative expression of representative trophoblast-associated genes in EpiSCs and iTSCs, highlighting upregulation of core trophoblast transcription factors and epithelial markers in iTSCs. (C) KEGG pathway enrichment analysis of genes upregulated in iTSCs compared with EpiSCs. Enriched pathways include signaling and structural processes associated with trophoblast identity, epithelial organization, and cell–matrix interactions.

### Embryoid body formation and blastocyst-like self-organization of pTSCs

The developmental capacity of pTSCs was evaluated by comparing their *in vitro* differentiation behavior with that of pEPSCs. During EB formation, the two cell types exhibited distinct morphologies. EBs derived from pEPSCs formed uniform, rounded spheroids with smooth outer surfaces and a relatively translucent internal appearance, consistent with the morphology of pluripotent stem cell aggregates (Fig. 6A). In contrast, EBs derived from pTSCs appeared darker, more compact, and heterogeneous, frequently displaying irregular or granular surface textures (Fig. 6A). Upon replating onto MEFs, EBs derived from pTSCs rapidly spread into flattened epithelial outgrowths. These outgrowths morphologically resembled trophoblast expansion observed during peri-implantation blastocyst development (Fig. 6B,C), supporting the acquisition of trophoblast-like properties. Building on recent advances in synthetic embryo systems in mouse and human models (28, 29), and emerging work in large animals (30-32), the potential for porcine EPSCs and TSCs to self-organize into blastocyst-like structures under three-dimensional, non-adherent culture conditions was assessed. Co-culture of pEPSCs with early-passage pTSCs maintained on MEF feeders resulted in the formation of compact epithelial vesicles resembling early blastocyst-like structures (Fig. 6D, left panel). In contrast, this self-organizing capacity was markedly reduced in long-term passaged pTSCs maintained under feeder-free conditions, which exhibited limited compaction and failed to form organized structures. Immunofluorescence analysis of successfully formed aggregates revealed polarized outer epithelial layers and epiblast-like structure, comparable to *ex vivo* blastocysts (Fig. 6E; Supplementary Fig. 5). However, despite these structural similarities, differences in lineage marker expression between blastocyst-like aggregates and native blastocysts were observed, indicating incomplete or imperfect lineage segregation. The efficiency of blastocyst-like structure formation was low, with organized aggregates arising in approximately 3-5% of co-cultured assemblies. Nevertheless, these findings demonstrate that early-passage, MEF-supported pTSCs retain a latent capacity to participate in the self-organization of embryonic structures. Collectively, these results establish a proof-of-concept framework for future efforts to reconstruct synthetic porcine embryos and to investigate extraembryonic lineage interactions in a large-animal context.

**Figure 6.**
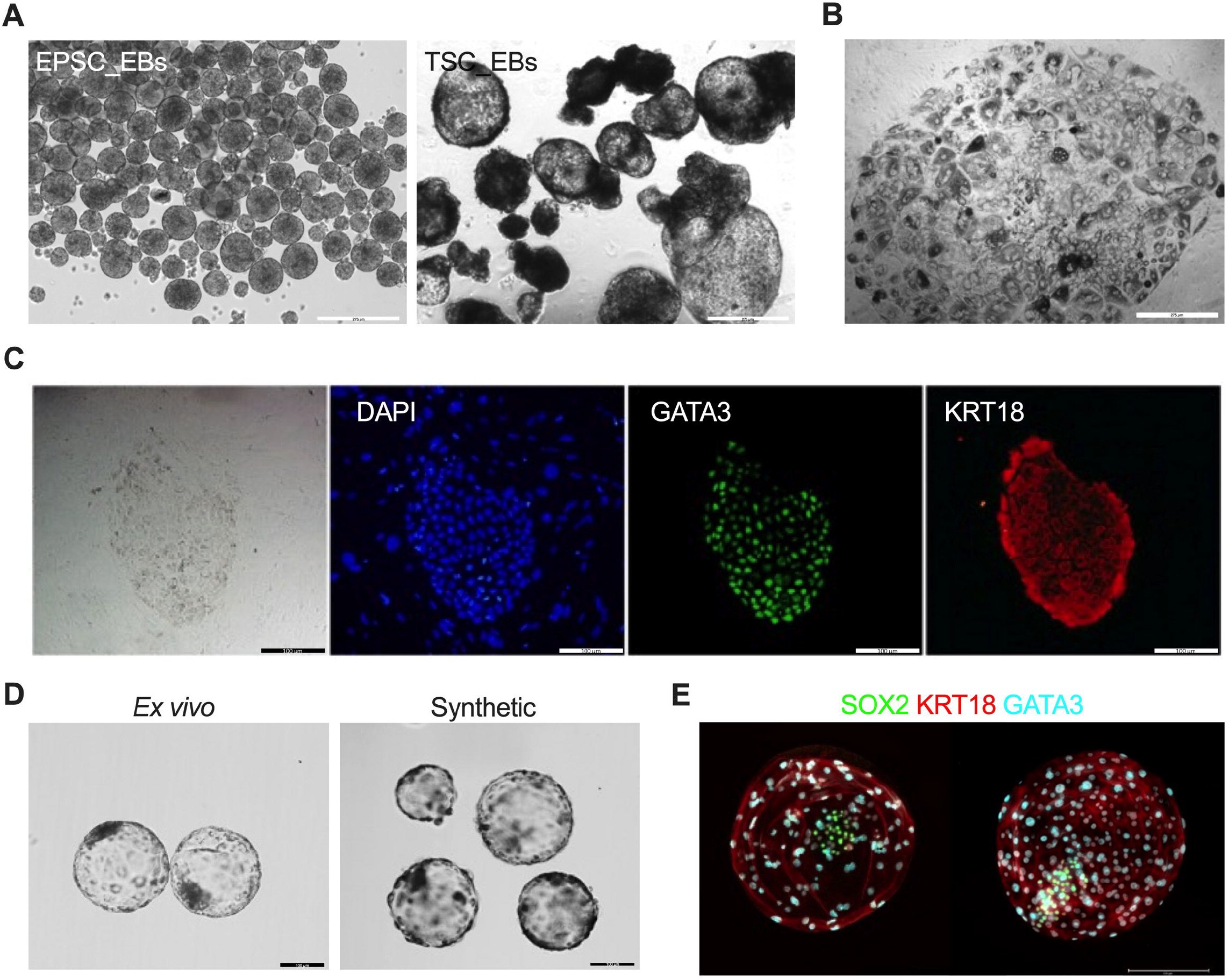
Embryoid body formation and blastocyst-like self-organization of pTSCs. (A) Representative bright-field images of embryoid bodies (EBs) generated from EPSCs (left) and pTSCs (right). EPSC-derived EBs form uniform, rounded spheroids with smooth surfaces, whereas pTSC-derived EBs appear darker, more compact, and morphologically heterogeneous. (B) Representative image of a pTSC-derived EB replated onto MEF feeders, showing rapid spreading as a flattened epithelial sheet resembling trophoblast outgrowth from porcine blastocysts. (C) Immunofluorescence analysis of replated pTSC-derived EB outgrowths showing expression of the trophoblast markers GATA3 and KRT18. (D) Comparison of *ex vivo* porcine blastocysts and *in vitro*-generated synthetic embryo-like structures formed by co-culture of pEPSCs and early-passage pTSCs. (E) Immunofluorescence analysis of *ex vivo* blastocysts and synthetic embryo–like structures showing spatial distribution of SOX2 (green), KRT18 (red), and GATA3 (cyan), highlighting partial architectural similarity between natural and synthetic structures. Scale bars, 100 μm.

### Lineage-restricted differentiation potential of pTSCs *in vivo*

To assess developmental potential *in vivo*, parental pEPSCs and induced pTSCs were transplanted subcutaneously into NOD-SCID mice. After six weeks, pTSC-derived grafts were significantly smaller (∼0.4 mm) than those formed by pEPSCs (∼1.2 mm) (Fig. 7A, B). Histological analysis indicated that pTSC-derived grafts contained a limited range of tissue types, including interstitial mesenchyme, adipose tissue, and muscle. In contrast, pEPSC-derived lesions formed classical teratomas with derivatives of all three germ layers, such as cartilage- and bone-like structures (Fig. 7C). Consistent with these observations, qPCR analysis demonstrated that pEPSC-derived tumors expressed markers of ectoderm (*NES, PAX6, ZIC1*), mesoderm (*DES, FGF8, T*), and endoderm (*HNF4A, GATA6, SOX17*). In contrast, pTSC-derived grafts lacked ectodermal and endodermal transcripts but maintained strong expression of mesodermal and extraembryonic markers, including *DES, FGF8*, and *T* (Fig. 7D). Collectively, these data indicate that induced pTSCs possess a lineage-restricted differentiation potential biased toward mesodermal and extraembryonic fates, consistent with the loss of pluripotency and acquisition of stable trophoblast identity.

**Figure 7.**
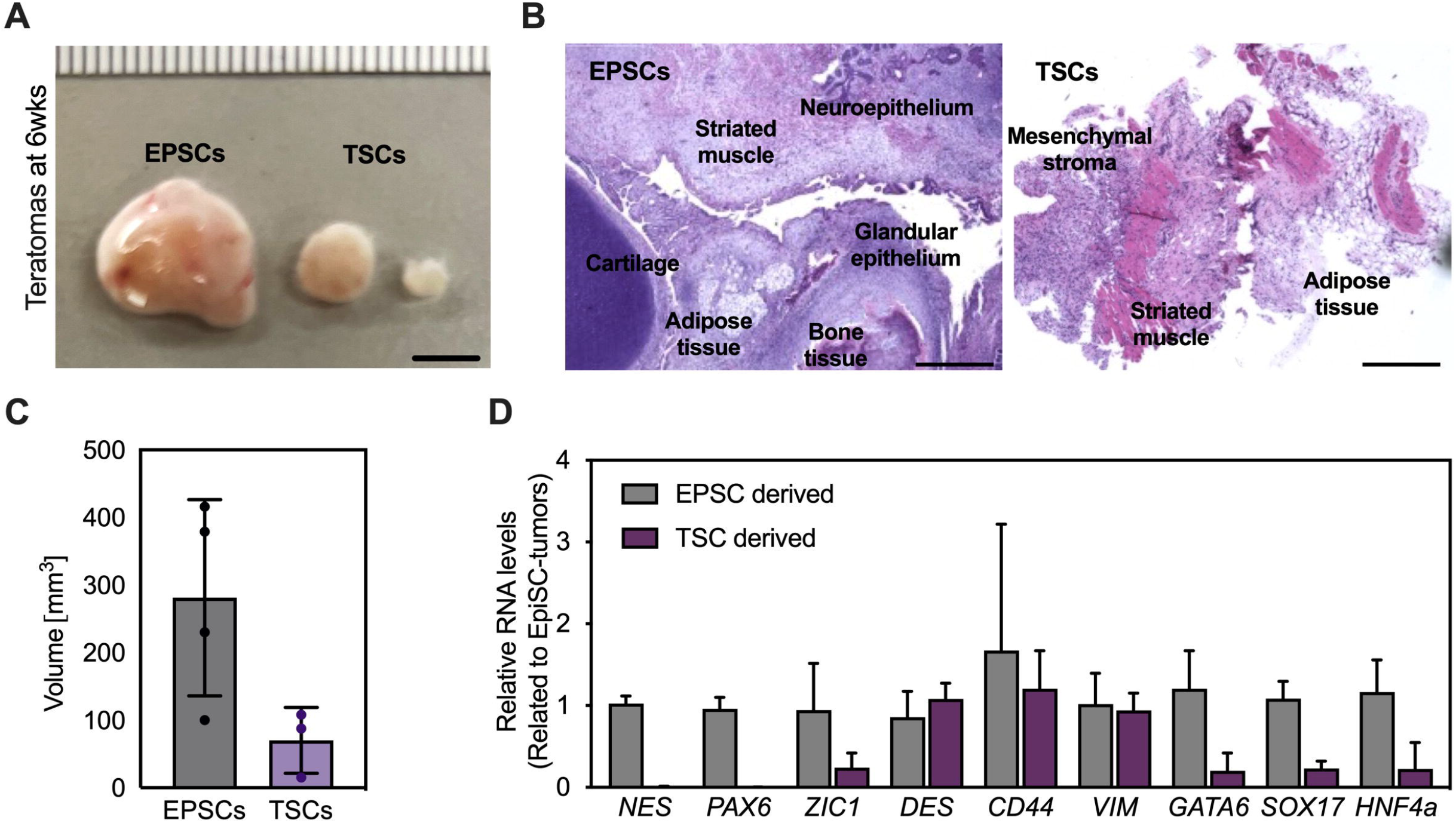
Lineage-restricted differentiation potential of pTSCs *in vivo*. (A) Gross appearance of tumors formed 6 weeks after subcutaneous injection of pEPSCs and pTSCs into NOD-SCID mice. TSC-derived grafts were visibly smaller. Scale bar: 1 mm. (B) Histological sections (H&E) of EPSC-derived teratomas displayed structures from all three germ layers (e.g., neuroepithelium, cartilage, bone, glandular epithelium), while TSC-derived lesions predominantly contained mesenchymal stromal tissue, adipose, and striated muscle. (C) Quantification of lesion volumes derived from pEPSCs and pTSCs six weeks after transplantation. (D) Relative expression of trilineage marker genes from tumor tissues, normalized to EPSC-derived grafts. Ectodermal (*NES, PAX6, ZIC1*) and endodermal (*GATA6, SOX17, HNF4A*) transcripts were diminished in TSC grafts, while mesodermal markers (*DES, CD44, VIM*) remained comparably expressed. Scale bars: B, D = 100 μm; C = 1 mm.

## Discussion

This study establishes a defined experimental framework for inducing and characterizing pTSCs from pEPSCs. Through targeted manipulation of signaling pathways, together with transcriptomic profiling and *in vivo* functional analyses, we demonstrate that combined inhibition of the TGF-β/Activin and MEK/ERK pathways is sufficient to drive stable conversion of the trophoblast lineage. In contrast, the composition of the basal medium critically influences induction efficiency and cellular viability. Collectively, these findings position the porcine system within a broader cross-species context, bridging insights from murine, human, and bovine TSC models. Initial experiments revealed that trophoblast induction conditions established for human pluripotent stem cells are not directly transferable to porcine cells. Specifically, the BAP formulation used for human ESC-derived TSCs (17, 33) failed to sustain epithelial integrity in pigs, primarily due to the cytotoxic effects of the FGFR inhibitor PD173074. In contrast, CTB medium (15) supported epithelialization and stable proliferation. The requirement for a DMEM/F12-based medium supplemented with KSR mirrors human TSC culture conditions but differs from murine systems, which rely on N2B27-based formulations (34). This divergence underscores fundamental metabolic and extracellular niche differences among mammalian trophoblast lineages (35). Furthermore, feeder-assisted induction improved survival and epithelial transition, consistent with previous reports that feeder-derived extracellular matrix components or paracrine factors stabilize early trophoblast fates (36).

Upregulation of *KRT7* and *GATA3*, along with repression of pluripotency and hypoblast markers, demonstrates that porcine trophoblast induction follows a transcriptional trajectory analogous to the TE program described in human and primate embryos (37). Inhibitor experiments identify dual TGF-β/Activin and ERK inhibition as the central driver of porcine trophoblast induction, reflecting a conserved requirement across species. In mice, FGF4 and Activin A maintain TSC self-renewal through ERK and TGF-β signaling (38), whereas in human systems, TGF-β/Activin inhibition combined with WNT and EGF signaling supports TSC maintenance (17, 35). In pigs, the data indicate that TGF-β inhibition is necessary to release pluripotency-associated repression, while sustained ERK inhibition prevents premature differentiation and promotes stable epithelial organization. BMP4 signaling has been implicated in trophoblast induction in human and primate PSCs (17, 23), yet its temporal role remains unresolved in pigs. In human ESCs, transient BMP exposure favors mesodermal differentiation, whereas prolonged activation promotes trophoblast fate (39). Clarifying whether porcine cells exhibit similar temporal sensitivity is essential for defining how BMP and TGF-β signaling interact to establish extraembryonic competence in large-animal systems. During conversion, pEPSCs rapidly downregulated pluripotency and hypoblast programs while upregulating canonical trophoblast regulators, including *CDX2, GATA3, TEAD4, HAND1*, and *ELF5*, consistent with conserved trophoblast specification networks across mammals (33, 40). Transient expression of *EOMES* suggests a brief overlap with mesodermal programs before full trophoblast stabilization, paralleling intermediate states observed during human PSC-to-TSC reprogramming (41).

Feeder dependence has long defined TSC systems, particularly in rodents, where MEF-derived FGF4 and TGF-β family ligands maintain the extraembryonic ectoderm state (42, 43). In contrast, human and primate TSCs are feeder-independent and rely on exogenous EGF, WNT activation, and TGF-β inhibition (17, 35). Our results indicate that porcine TSCs occupy an intermediate configuration between these paradigms. On feeders, induced pTSCs maintained higher *CDX2* expression and compact epithelial morphology, resembling early ExE-like progenitors. Upon transition to Matrigel, cells flattened, proliferated more rapidly, and downregulated *CDX2*, thereby adopting features more closely resembling those of human CTB-like cells. This feeder-dependent dichotomy likely reflects differential exposure to paracrine FGF and TGF-β signaling from MEFs versus intrinsic WNT-driven, TGF-β-repressed signaling in feeder-free conditions. These observations support a model in which trophoblast stemness exists along a continuum of epithelial states modulated by extracellular signaling environments rather than as a single fixed identity.

Pathway-level transcriptomic analysis further validated lineage conversion, revealing enrichment of extracellular matrix organization, epithelial junctions, and hormone-related pathways, accompanied by silencing of pluripotency and hypoblast programs. These features align with hallmarks of early TE and extraembryonic ectoderm in other species(44, 45). The divergence in KEGG pathway enrichment between EPSCs and pTSCs reflects the functional demands of each lineage: EPSCs exhibit enrichment of WNT, TGF-β, and MAPK signaling pathways associated with epiblast maintenance, whereas pTSCs show activation of PI3K-Akt signaling, adhesion, vesicle trafficking, and metabolic pathways characteristic of trophoblast function. These transcriptional changes provide strong molecular evidence for directed trophoblast differentiation and underscore the value of pathway-level analyses for assessing lineage fidelity. Functional validation using teratoma assays further demonstrated lineage restriction. While pEPSCs generated classical multi-lineage teratomas containing ectodermal, mesodermal, and endodermal derivatives, induced pTSCs formed small, non-teratomatous lesions expressing only mesodermal and extraembryonic markers. This restricted differentiation potential mirrors the behavior of *bona fide* human and bovine TSCs in xenograft assays (24, 35), thereby reinforcing the trophoblast identity of induced pTSCs. Thus, the porcine system displays functional features consistent with trophoblast identity. Together, our data demonstrate that porcine EPSCs can be directly induced into a stable trophoblast-like state under defined signaling conditions. The porcine system occupies an evolutionary position between rodent and primate models; therefore, its establishment expands the comparative framework for studying conserved and divergent mechanisms of trophoblast specification (46). Further refinement of the process including temporal modulation of BMP4, long-term epigenetic profiling, and direct comparison with embryo-derived TE, will be essential to validate lineage fidelity. Together, this platform provides a valuable large-animal model for dissecting trophoblast-specific signaling, implantation biology, and placental pathophysiology.

## Conclusion

In summary, we established a reproducible, chemically defined protocol for inducing TSCs from pESCs. Our findings identify TGF-β/Activin inhibition as necessary and MEK/ERK blockade as sufficient for robust trophoblast conversion, revealing a signaling logic conserved in primate systems but distinct from that in rodent models. Transcriptomic and *in vivo* analyses demonstrate that induced pTSCs acquire stable epithelial identity and lineage-restricted developmental potential consistent with trophoblast fate. This platform provides a powerful large-animal model for dissecting conserved and species-specific mechanisms of trophoblast specification, implantation biology, and placental pathophysiology.

## Materials and Methods

### Ethics statement

All experiments involving live animals were performed as per the approved guidelines of the University of Missouri, Institutional Animal Care and Use Committee protocol # 63641.

### EPSC culture

pEPSC were derived and cultured in ‘3iLA’ medium. Briefly, 3iLA is a N2B27-based medium supplemented with 0.3□µM CHIR99021 (Tocris), 0.3□µM WH-4-023 (Tocris), 5□µM IWR-endo-1 (Tocris,), 50□µg□ml^−1^ L-ascorbic acid 2-phosphate, a Rho-associated kinase (ROCK) inhibitor, 5 µM Y-27632 (72304, STEMCELL Technologies), 10□ng□ml^−1^ recombinant human LIF (PeproTech), 20□ng□ml^−1^ Activin A (PeproTech), 10 ng ml^−1^ bFGF (PeproTech). A basal N2B27 medium contains a 1:1 mixture of DMEM/F-12 (21331-020, Gibco) and Neurobasal A (10888-022, Thermo Fisher Scientific) supplemented with 0.5% v/v N2 supplement (17502048, Thermo Fisher Scientific), 1% v/v B27 supplement (10889–038, Thermo Fisher Scientific), 5% v/v KnockOut Serum Replacement (KSR, Thermo Fisher Scientific), 1% penicillin-streptomycin (PS; 15140122, Thermo Fisher Scientific), GlutaMAX (35050061, Thermo Fisher Scientific), non-essential amino acids (NEAA, 35050061, Thermo Fisher Scientific), and β-mercaptoethanol (β-ME; 31350-010, Thermo Fisher Scientific).

To derive the primary outgrowths, the expanded blastocysts at day 7 were seeded onto inactivated CF-1 mouse embryonic fibroblasts (MEFs), which were plated at 3 × 10^5^ cells per cm^2^ onto a gelatinized 4-well plate (Nunc) in 3iLAF medium. Under these conditions, the blastocysts attached, and primary outgrowths developed within 48□h. The media was changed every other day. After 7-9 days, individual epiblast colonies were passaged onto feeders in 4-well plates (p1) and thereafter in 6-well culture plates (p2). All cell lines were passaged as clumps by dissociation with a 1:1 mix of Accutase (A1110501, ThermoFisher Scientific) and TrypLE Express (Gibco) in the presence of 10 µM Y-27632, which was removed after 24 hr. EPSCs were passaged every 4 days, and the medium was changed daily. AP staining of EPSCs was performed using the Alkaline Phosphatase Detection Kit (00-0055, Stemgent), according to the kit’s instructions. Cell Line Genetics LLC performed karyotypic analysis. All cell cultures in this paper were maintained in a humidified incubator set at 38.5 □C, 5% CO_2_ unless stated otherwise.

### TSC derivation from PSCs or embryos and their maintenance

EPSCs were induced to TE over the course of 5 days by plating PSCs at passage 60-65 onto MEFs in inductive medium, CT (47). In detail, pEPSCs (2 × 10^5^ per well of 6-well plates) were seeded onto MEFs and cultured in 3iLA medium. The following day (day 0), the culture medium was replaced with TS medium for pTSC derivation. A basal TS medium contains advanced DMEM/F12 (Gibco) supplemented with 5% KOSR, 0.3% w/v bovine serum albumin (BSA), 1% sodium-pyruvate 1% non-essential amino acids, 1% GlutaMAX (ThermoFisher 35050061), 0.1 mM β-ME, 1% PS, 1.25 mM N-Acetyl-L-cysteine, 10 mM Nicotinamide, 10□µg□ml^−1^ L-ascorbic acid 2-phosphate, 1 μM A83-01 or 2 μM SB431542 (Selleck), 2 μM CHIR99021 (Selleck), 20 ng ml^−1^ hrEGF (RD), 20 nM trichostatin A (TSA) and 5 μM Y-27632. After 5 days, the TE-like cells passaged at a 1:3 ratio with TrypLE and cultured on the Matrigel-coated 6-well plate in iTE medium continuously for a week. For testing “BAP” conditions we used N2B27 medium supplemented with 20 ng ml^−1^ BMP4 (RD), 0.5 μM A83-01 and 1 μM PD0325901 or 0.1 µM PD173074.

### Engraftment of TSCs into NOD-SCID mice

Induced or derived EpiSC or TSC (5 × 10^6^ cells) were resuspended in 100 μL of DMEM-Matrigel solution (1:1) supplemented with 10 µM Y-27632 and were injected subcutaneously into 6-week-old NOD/SCID/γc^−/−^ mice (NSG mice; The Jackson Laboratory). After 5-6 weeks, the volume of dissected lesions (VL) was calculated by the ellipsoidal formula VL = 0.5 (length × width^2^). Then, the lesions were fixed in 10% formalin, paraffin-embedded, and sectioned for Hematoxylin and eosin staining and immunostaining.

### RNA analysis and qPCR

Total RNA was isolated using the RNeasy® Total RNA Isolation Mini Kit (Qiagen, Life Technologies) following the manufacturer’s protocol, and complementary DNA (cDNA) was synthesized using the cDNA Synthesis Kit (Applied Biosystems). Quantitative real-time polymerase chain reaction (qPCR) analysis was performed using the Power SYBR Green PCR Master Mix with the ABI7500 Real-Time PCR System (Applied Biosystems). Target gene expression was normalized to the housekeeping genes RN18S and ACTB for quantification. The primer sequences used for amplification are listed in Supplementary Table 1.

### Bulk RNA-seq and analysis

RNA samples were sent to Azenta US, Inc. (South Plainfield, NJ) for bulk RNA sequencing. FASTQ files from 20 million reads per sample were aligned to the *Sus scrofa* reference genome (Sscrofa11.1/susScr11) using the STAR aligner. Using DESeq2, a comparison of gene expression between EPSC and iTSC was performed with an adjusted p-value < 0.05 as the cutoff for statistical significance. The package clusterProfiler (version 3.0.4) was used for GO enrichment analysis.

## Data availability

A total of 18 transcriptomic data sets generated for this article can be accessed via NCBI Sequence Read Archive (SRA) SUB15985520.

## Immunofluorescence

Early embryos were rinsed three times with 0.1% PBS/polyvinylpyrrolidone, fixed in 4% paraformaldehyde in PBS for 5 min, and permeabilized in 0.5% Triton X-100/PBS for 30 min. Fixed samples were then blocked for 2 h in a buffer containing 5% bovine serum albumin and 0.1% Triton X-100 in PBS. Samples were incubated with primary antibodies overnight at 4°C, and embryos were then treated with secondary antibodies for 1 h. Nuclear DNA was counterstained by DAPI (Thermo Fisher Scientific) for 5 min. Samples were mounted and photographed by using a Leica DM 4000B microscope using the Leica Application Suite Advanced Fluorescence software (Leica Microsystems, Wetzlar, Germany). All antibody information is shown in the Supplementary Table 2.

## Author Contributions

C.P. and Y.J. conceived the project and designed the experiments; C.P. and Y.J. performed derivation of the cell lines, differentiation. C.P., Y.J., and J.W. performed characterization of the lines; C.P. generated the libraries and performed transcriptomic analysis; C.P. wrote the initial draft; B.P.T. revised the manuscript based on the input from all authors. All authors approved the final draft for submission.

## Guarantor Statement

B.P.T. (Corresponding author) is the guarantor of this work and, as such, had full access to all the data in the study and takes responsibility for the integrity of the data, the accuracy of the analyses, and the decision to submit the manuscript for publication.

## Competing Interest Statement

All authors declare no competing interests or conflicts of interest.

## Acknowledgements

This research was supported by funding from Genus Plc and University of Missouri.

## Supplementary Information

This manuscript contains Supplementary information.

## Notes

### Competing Interest Statement

The authors have declared no competing interest.

## References

1. Hay, W. W., Jr. (1994) Placental transport of nutrients to the fetus Horm Res 42, 215–222

2. Megli, C. J., and Coyne, C. B. (2022) Infections at the maternal-fetal interface: an overview of pathogenesis and defence Nat Rev Microbiol 20, 67–82

3. Tal, R., and Taylor, H. S. (2000) Endocrinology of Pregnancy, MDText.com, Inc., South Dartmouth (MA),

4. Roberts, R. M., Green, J. A., and Schulz, L. C. (2016) The evolution of the placenta Reproduction 152, R179–189

5. Moffett, A., and Loke, C. (2006) Immunology of placentation in eutherian mammals Nat Rev Immunol 6, 584–594

6. Friess, A. E., Sinowatz, F., Skolek-Winnisch, R., and Träutner, W. (1980) The placenta of the pig: I. Finestructural changes of the placental barrier during pregnancy Anatomy and embryology 158, 179–191

7. Bazer, F. W., and Johnson, G. A. (2014) Pig blastocyst-uterine interactions Differentiation 87, 52–65

8. Liu, Y., Fan, X., Wang, R., Lu, X., Dang, Y.-L., Wang, H. et al.. (2018) Single-cell RNA-seq reveals the diversity of trophoblast subtypes and patterns of differentiation in the human placenta Cell research 28, 819–832

9. Robbins, J. R., Skrzypczynska, K. M., Zeldovich, V. B., Kapidzic, M., and Bakardjiev, A. I. (2010) Placental syncytiotrophoblast constitutes a major barrier to vertical transmission of Listeria monocytogenes PLoS Pathog 6, e1000732

10. Wallace, A. E., Fraser, R., and Cartwright, J. E. (2012) Extravillous trophoblast and decidual natural killer cells: a remodelling partnership Human reproduction update 18, 458–471

11. Friess, A. E., Sinowatz, F., Skolek-Winnisch, R., and Träutner, W. (1981) The placenta of the pig: II. The ultrastructure of the areolae Anatomy and Embryology 163, 43–53

12. McLendon, B. A., Kramer, A. C., Seo, H., Burghardt, R. C., Bazer, F. W., Wu, G. et al.. (2022) Temporal and spatial expression of aquaporins 1, 5, 8, and 9: potential transport of water across the endometrium and chorioallantois of pigs Placenta 124, 28–36

13. Suleman, M., Malgarin, C. M., Detmer, S. E., Harding, J. C. S., and MacPhee, D. J. (2019) The porcine trophoblast cell line PTr2 is susceptible to porcine reproductive and respiratory syndrome virus-2 infection Placenta 88, 44–51

14. Hou, D., Su, M., Li, X., Li, Z., Yun, T., Zhao, Y. et al.. (2015) The Efficient Derivation of Trophoblast Cells from Porcine In Vitro Fertilized and Parthenogenetic Blastocysts and Culture with ROCK Inhibitor Y-27632 PLoS One 10, e0142442

15. Okae, H., Toh, H., Sato, T., Hiura, H., Takahashi, S., Shirane, K. et al.. (2018) Derivation of Human Trophoblast Stem Cells Cell Stem Cell 22, 50–63 e56

16. Guo, G., Stirparo, G. G., Strawbridge, S. E., Spindlow, D., Yang, J., Clarke, J. et al.. (2021) Human naive epiblast cells possess unrestricted lineage potential Cell Stem Cell 28, 1040–1056 e1046

17. Viukov, S., Shani, T., Bayerl, J., Aguilera-Castrejon, A., Oldak, B., Sheban, D. et al.. (2022) Human primed and naive PSCs are both able to differentiate into trophoblast stem cells Stem Cell Reports 17, 2484–2500

18. Zhi, M., Zhang, J., Tang, Q., Yu, D., Gao, S., Gao, D. et al.. (2022) Generation and characterization of stable pig pregastrulation epiblast stem cell lines Cell Res 32, 383–400

19. Choi, K. H., Lee, D. K., Kim, S. W., Woo, S. H., Kim, D. Y., and Lee, C. K. (2019) Chemically Defined Media Can Maintain Pig Pluripotency Network In Vitro Stem Cell Reports 13, 221–234

20. Kinoshita, M., Kobayashi, T., Planells, B., Klisch, D., Spindlow, D., Masaki, H. et al.. (2021) Pluripotent stem cells related to embryonic disc exhibit common self-renewal requirements in diverse livestock species Development 148,

21. Ruan, D., Xuan, Y., Tam, T., Li, Z., Wang, X., Xu, S. et al.. (2024) An optimized culture system for efficient derivation of porcine expanded potential stem cells from preimplantation embryos and by reprogramming somatic cells Nat Protoc 19, 1710–1749

22. Xu, J., Yu, L., Guo, J., Xiang, J., Zheng, Z., Gao, D. et al.. (2019) Generation of pig induced pluripotent stem cells using an extended pluripotent stem cell culture system Stem Cell Res Ther 10, 193

23. Xu, R. H., Chen, X., Li, D. S., Li, R., Addicks, G. C., Glennon, C. et al.. (2002) BMP4 initiates human embryonic stem cell differentiation to trophoblast Nat Biotechnol 20, 1261–1264

24. Wang, Y., Ming, H., Yu, L., Li, J., Zhu, L., Sun, H. X. et al.. (2023) Establishment of bovine trophoblast stem cells Cell Rep 42, 112439

25. Saha, B., Ganguly, A., Home, P., Bhattacharya, B., Ray, S., Ghosh, A. et al.. (2020) TEAD4 ensures postimplantation development by promoting trophoblast self-renewal: An implication in early human pregnancy loss Proc Natl Acad Sci U S A 117, 17864–17875

26. Kubaczka, C., Senner, C. E., Cierlitza, M., Arauzo-Bravo, M. J., Kuckenberg, P., Peitz, M. et al.. (2015) Direct Induction of Trophoblast Stem Cells from Murine Fibroblasts Cell Stem Cell 17, 557–568

27. Park, C. H., Jeoung, Y. H., Uh, K. J., Park, K. E., Bridge, J., Powell, A. et al.. (2021) Extraembryonic Endoderm (XEN) Cells Capable of Contributing to Embryonic Chimeras Established from Pig Embryos Stem Cell Reports 16, 212–223

28. Rivron, N. C., Frias-Aldeguer, J., Vrij, E. J., Boisset, J. C., Korving, J., Vivie, J. et al.. (2018) Blastocyst-like structures generated solely from stem cells Nature 557, 106–111

29. Rossant, J., and Tam, P. P. L. (2022) Early human embryonic development: Blastocyst formation to gastrulation Dev Cell 57, 152–165

30. Pinzon-Arteaga, C. A., Wang, Y., Wei, Y., Ribeiro Orsi, A. E., Li, L., Scatolin, G. et al.. (2023) Bovine blastocyst-like structures derived from stem cell cultures Cell Stem Cell 30, 611–616 e617

31. Kim, H., and Kim, E. (2024) Current Status of Synthetic Mammalian Embryo Models Int J Mol Sci 25,

32. Xiang, J., Wang, H., Shi, B., Li, J., Liu, D., Wang, K. et al.. (2024) Pig blastocyst-like structure models from embryonic stem cells Cell Discov 10, 72

33. Amita, M., Adachi, K., Alexenko, A. P., Sinha, S., Schust, D. J., Schulz, L. C. et al.. (2013) Complete and unidirectional conversion of human embryonic stem cells to trophoblast by BMP4 Proc Natl Acad Sci U S A 110, E1212–1221

34. Briseno, C. G., Satpathy, A. T., Davidson, J. T. t., Ferris, S. T., Durai, V., Bagadia, P. et al.. (2018) Notch2-dependent DC2s mediate splenic germinal center responses Proc Natl Acad Sci U S A 115, 10726–10731

35. Soncin, F., Morey, R., Bui, T., Requena, D. F., Cheung, V. C., Kallol, S. et al.. (2022) Derivation of functional trophoblast stem cells from primed human pluripotent stem cells Stem Cell Reports 17, 1303–1317

36. Kubaczka, C., Senner, C., Arauzo-Bravo, M. J., Sharma, N., Kuckenberg, P., Becker, A. et al.. (2014) Derivation and maintenance of murine trophoblast stem cells under defined conditions Stem Cell Reports 2, 232–242

37. Blakeley, P., Fogarty, N. M., del Valle, I., Wamaitha, S. E., Hu, T. X., Elder, K. et al.. (2015) Defining the three cell lineages of the human blastocyst by single-cell RNA-seq Development 142, 3151–3165

38. Tanaka, S., Kunath, T., Hadjantonakis, A. K., Nagy, A., and Rossant, J. (1998) Promotion of trophoblast stem cell proliferation by FGF4 Science 282, 2072–2075

39. Bernardo, A. S., Faial, T., Gardner, L., Niakan, K. K., Ortmann, D., Senner, C. E. et al.. (2011) BRACHYURY and CDX2 mediate BMP-induced differentiation of human and mouse pluripotent stem cells into embryonic and extraembryonic lineages Cell Stem Cell 9, 144–155

40. Roberts, R. M., and Fisher, S. J. (2011) Trophoblast stem cells Biol Reprod 84, 412–421

41. Dong, C., Beltcheva, M., Gontarz, P., Zhang, B., Popli, P., Fischer, L. A. et al.. (2020) Derivation of trophoblast stem cells from naive human pluripotent stem cells Elife 9,

42. Yang, W., Klaman, L. D., Chen, B., Araki, T., Harada, H., Thomas, S. M. et al.. (2006) An Shp2/SFK/Ras/Erk signaling pathway controls trophoblast stem cell survival Dev Cell 10, 317–327

43. Uy, G. D., Downs, K. M., and Gardner, R. L. (2002) Inhibition of trophoblast stem cell potential in chorionic ectoderm coincides with occlusion of the ectoplacental cavity in the mouse Development 129, 3913–3924

44. Simmons, D. G., and Cross, J. C. (2005) Determinants of trophoblast lineage and cell subtype specification in the mouse placenta Dev Biol 284, 12–24

45. Au, H. K., Peng, S. W., Guo, C. L., Lin, C. C., Wang, Y. L., Kuo, Y. C. et al.. (2021) Niche Laminin and IGF-1 Additively Coordinate the Maintenance of Oct-4 Through CD49f/IGF-1R-Hif-2alpha Feedforward Loop in Mouse Germline Stem Cells Front Cell Dev Biol 9, 646644

46. Sah, N., and Soncin, F. (2024) Conserved and divergent features of trophoblast stem cells J Mol Endocrinol 72,

47. Wei, Y., Wang, T., Ma, L., Zhang, Y., Zhao, Y., Lye, K. et al.. (2021) Efficient derivation of human trophoblast stem cells from primed pluripotent stem cells Sci Adv 7,

