## Supplementary Information for "Directed conversion of porcine extended pluripotent stem cells into trophoblast-like stem cells through modulation of conserved TGF-β and ERK signaling pathways"

**This PDF file includes:**

Supplementary Figures
- sFigure 1. Characterization of porcine blastocyst outgrowth and evaluation of basal media during early trophoblast induction.
- sFigure 2. Effects of signaling inhibition on pluripotency and hypoblast-associated gene expression during early trophoblast induction.
- sFigure 3. Dynamic regulation of hypoblast-associated markers during early trophoblast induction.
- sFigure 4. Differential gene expression and pathway enrichment analysis between EpiSCs and TSCs.
- sFigure 5. Comparative marker expression in *ex vivo* blastocysts and synthetic embryo-like structures

Supplementary Tables
- sTable 1. RT-qPCR primers
- sTable 2. Antibody information

### Supplementary Figures

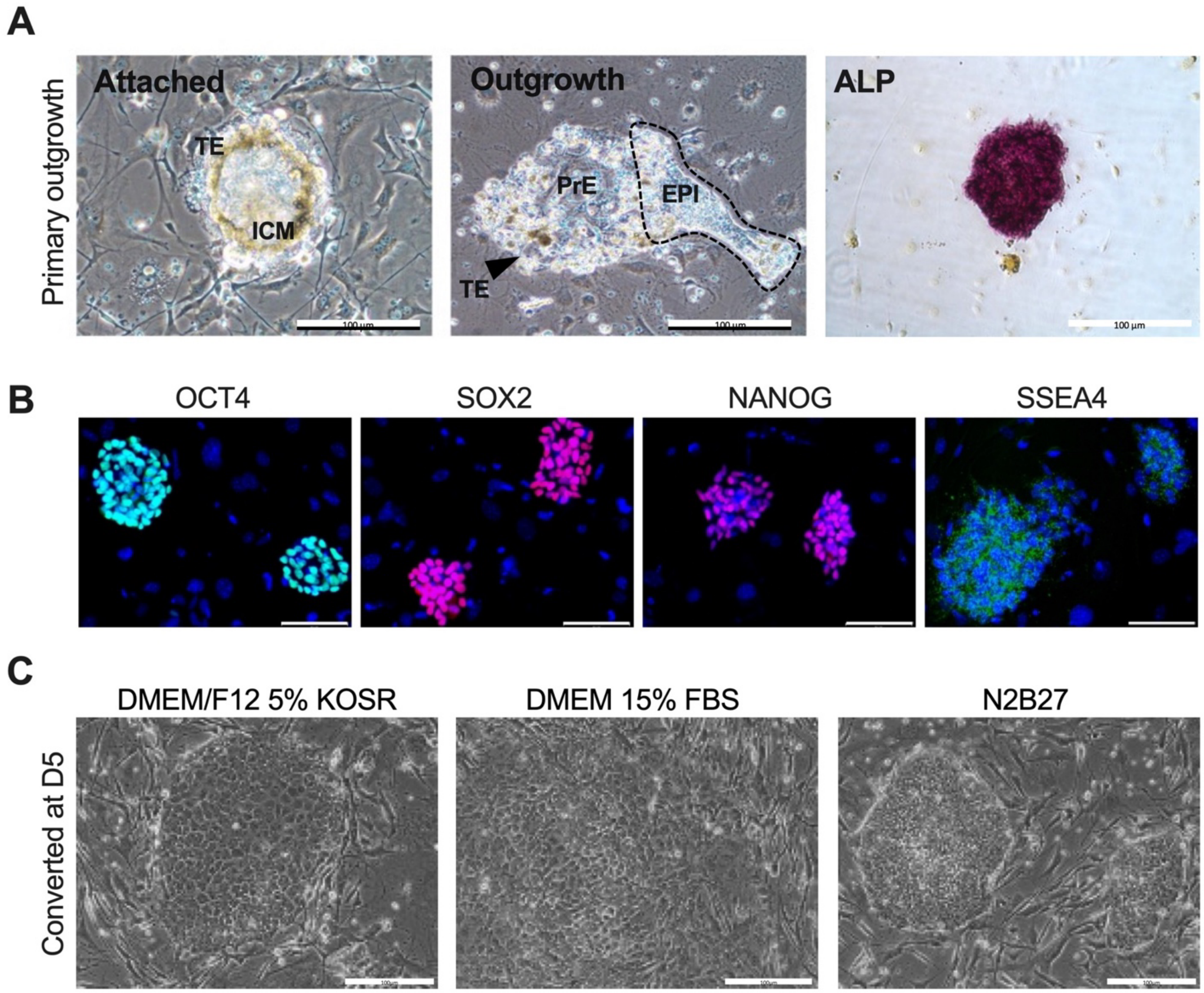


**Supplementary Figure 1. Characterization of porcine blastocyst outgrowth and evaluation of basal media during early trophoblast induction.** Related to Figure 1

(A) Representative images of porcine blastocyst outgrowth during early attachment and lineage segregation. At day 2 post-attachment, primary outgrowths display an attached blastocyst with identifiable trophectoderm (TE) and inner cell mass (ICM). By day 3, expanded outgrowths contain spatially segregated lineages, including epiblast (EPI; outlined by dotted line), primitive endoderm (PrE), and surrounding TE cells. Epiblast colonies isolated at day 4-5 exhibit strong alkaline phosphatase (ALP) activity, consistent with pluripotent identity.

(B) Immunofluorescence staining of EPSCs showing robust expression of the pluripotency markers OCT4, SOX2, NANOG, and SSEA4. Scale bars, 75 µm.

(C) Morphology of cells at day 5 of TSC induction under three different basal media conditions: DMEM/F12 supplemented with 5% KSR, DMEM/F12 supplemented with 15% FBS, and N2B27. DMEM/12 conditions support epithelial colony formation, with differences in colony compactness and surrounding cell morphology.
Scale bars, 100 μm**.**

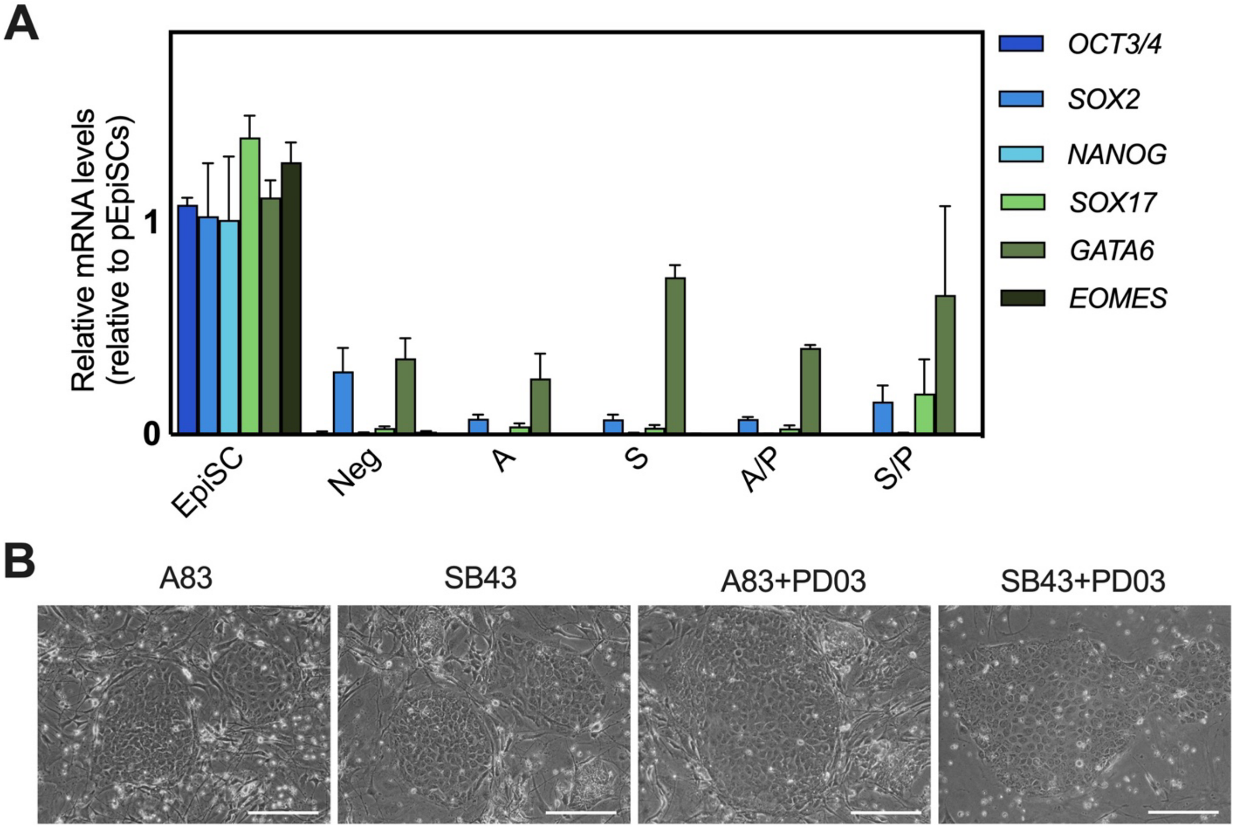


**Supplementary Figure 2. Effects of signaling inhibition on pluripotency and hypoblast-associated gene expression during early trophoblast induction.** Related to Figure 2

**(**A) Quantitative RT-PCR analysis of pluripotency-associated (*OCT3/4, SOX2, NANOG*) and hypoblast-associated (*SOX17, GATA6, EOMES*) gene expression in porcine EpiSCs and cells subjected to trophoblast induction under the inhibitor conditions shown in Figure 2. Expression levels are shown relative to parental EPSCs. Inhibition of the TGF-β/Activin pathway, alone or in combination with MEK/ERK inhibition, results in efficient suppression of pluripotency markers and variable regulation of hypoblast-associated genes, including partial re-expression of *GATA6* under specific conditions.

(B) Representative phase-contrast images of induced cultures at day 5 under A83-01, SB431542, A83-01 + PD0325901, and SB431542 + PD0325901 conditions, illustrating differences in epithelial colony morphology and compaction during early trophoblast induction. Scale bars, 100 µm.

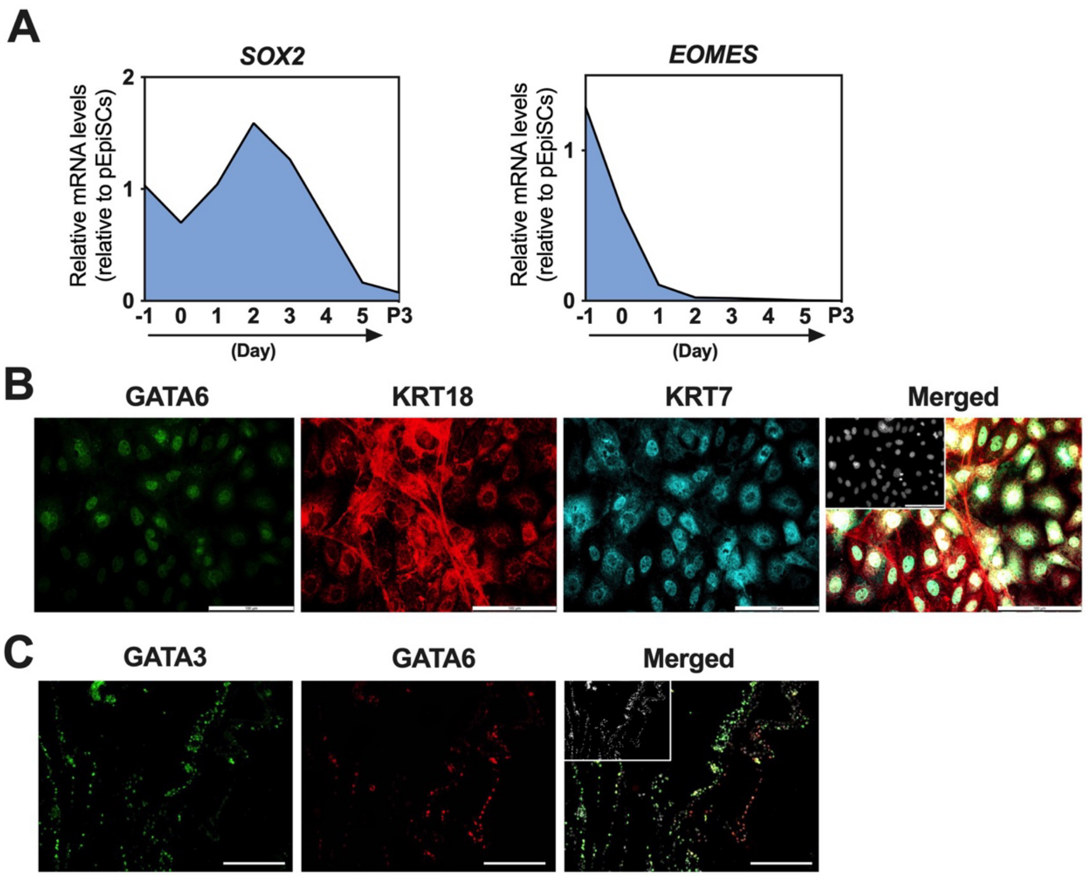


**Supplementary Figure 3. Dynamic regulation of hypoblast-associated markers during early trophoblast induction.** Related to Figure 3

(A) Quantitative RT-PCR analysis of *SOX2* and *EOMES* expression during early induction of porcine EPSCs under optimized trophoblast-inducing conditions. Expression levels are shown relative to parental EPSCs.

(B) Immunofluorescence analysis of induced TSC-like cells demonstrating detectable GATA6 protein in a subset of cells. Cells were co-stained for trophoblast markers to assess lineage context. Scale bars, 100 μm.

(C) Immunofluorescence analysis of extraembryonic tissues of day-25 porcine conceptuses. Scale bars, 250 μm.

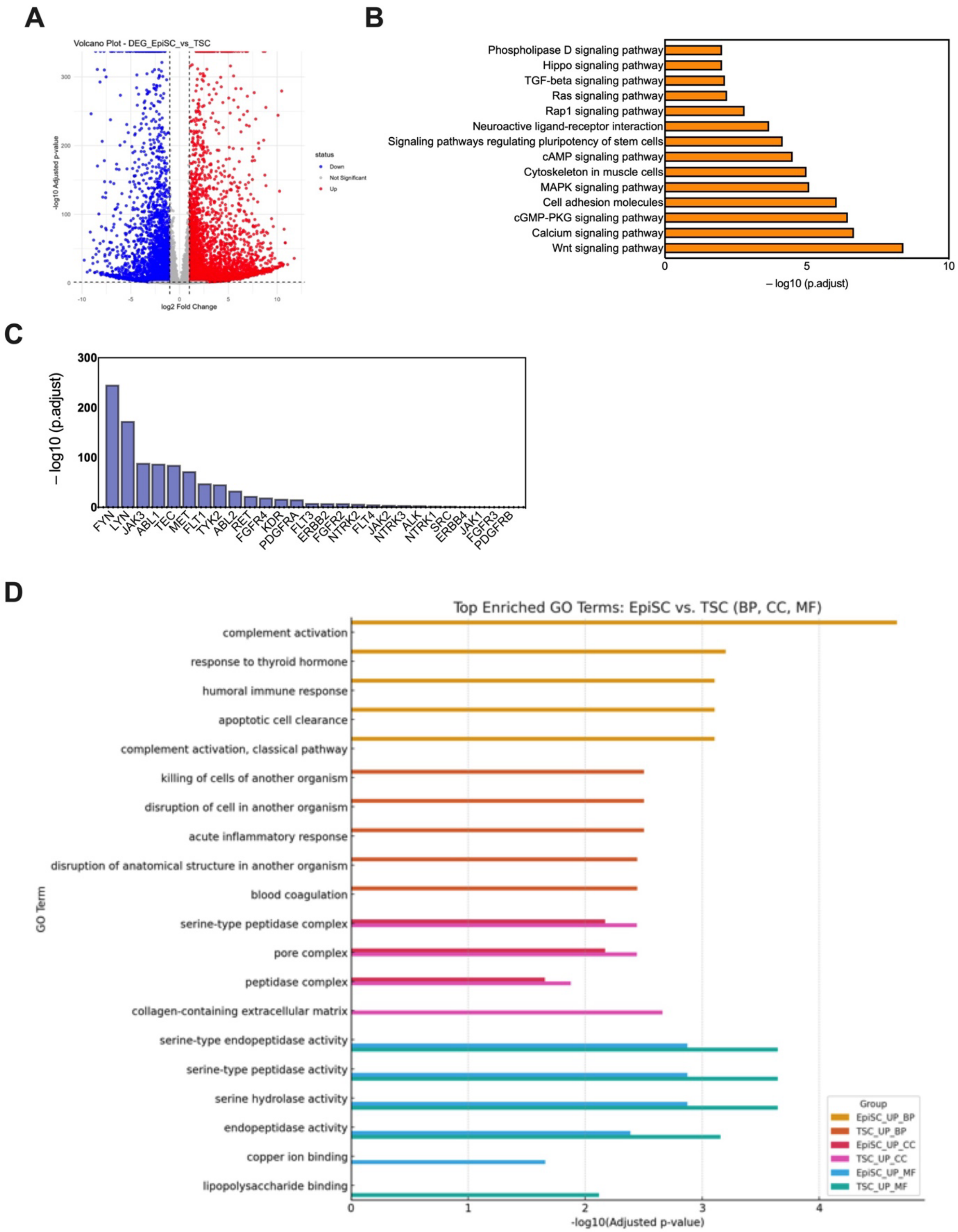

A
Volcano Plot - DEG_EpiSC_vs_TSC
-log10 Adjusted p-value
log2 Fold Change
status
Down
Not Significant
Up
B
Phospholipase D signaling pathway
Hippo signaling pathway
TGF-beta signaling pathway
Ras signaling pathway
Rap1 signaling pathway
Neuroactive ligand-receptor interaction
Signaling pathways regulating pluripotency of stem cells
cAMP signaling pathway
Cytoskeleton in muscle cells
MAPK signaling pathway
Cell adhesion molecules
cGMP-PKG signaling pathway
Calcium signaling pathway
Wnt signaling pathway
– log10 (p.adjust)
C
– log10 (p.adjust)
FYN
LYN
JAK3
ABL1
TEC
MET
FLT1
TYK2
ABL2
RET
FGFR4
KDR
PDGFRA
FLT3
ERBB2
FGFR2
NTRK2
FLT4
JAK2
NTRK3
ALK
NTRK1
SRC
ERBB4
JAK1
FGFR3
PDGFRB
D
Top Enriched GO Terms: EpiSC vs. TSC (BP, CC, MF)
complement activation
response to thyroid hormone
humoral immune response
apoptotic cell clearance
complement activation, classical pathway
killing of cells of another organism
disruption of cell in another organism
acute inflammatory response
disruption of anatomical structure in another organism
blood coagulation
serine-type peptidase complex
pore complex
peptidase complex
collagen-containing extracellular matrix
serine-type endopeptidase activity
serine-type peptidase activity
serine hydrolase activity
endopeptidase activity
copper ion binding
lipopolysaccharide binding
GO Term
-log10(Adjusted p-value)
Group
EpiSC_UP_BP
TSC_UP_BP
EpiSC_UP_CC
TSC_UP_CC
EpiSC_UP_MF
TSC_UP_MF

**Supplementary Figure 4. Differential gene expression and pathway enrichment analysis between EPSCs and TSCs.** Related to Figure 5

(A) Volcano plot showing differentially expressed genes between EPSCs and TSCs. Upregulated and downregulated genes are indicated, with significance defined by adjusted p-value and fold-change thresholds.

(B) KEGG pathway enrichment analysis of genes upregulated in EPSCs relative to TSCs, highlighting signaling pathways associated with pluripotency and epiblast identity.

(C) Enrichment of tyrosine kinase (TK)- associated genes among differentially expressed genes between EPSCs and TSCs, ranked by statistical significance.

(D) Gene Ontology (GO) enrichment analysis of differentially expressed genes between EPSCs and TSCs across Biological Process (BP), Cellular Component (CC), and Molecular Function (MF) categories. Bars represent the most significantly enriched terms, shown as −log10 (adjusted p-value).

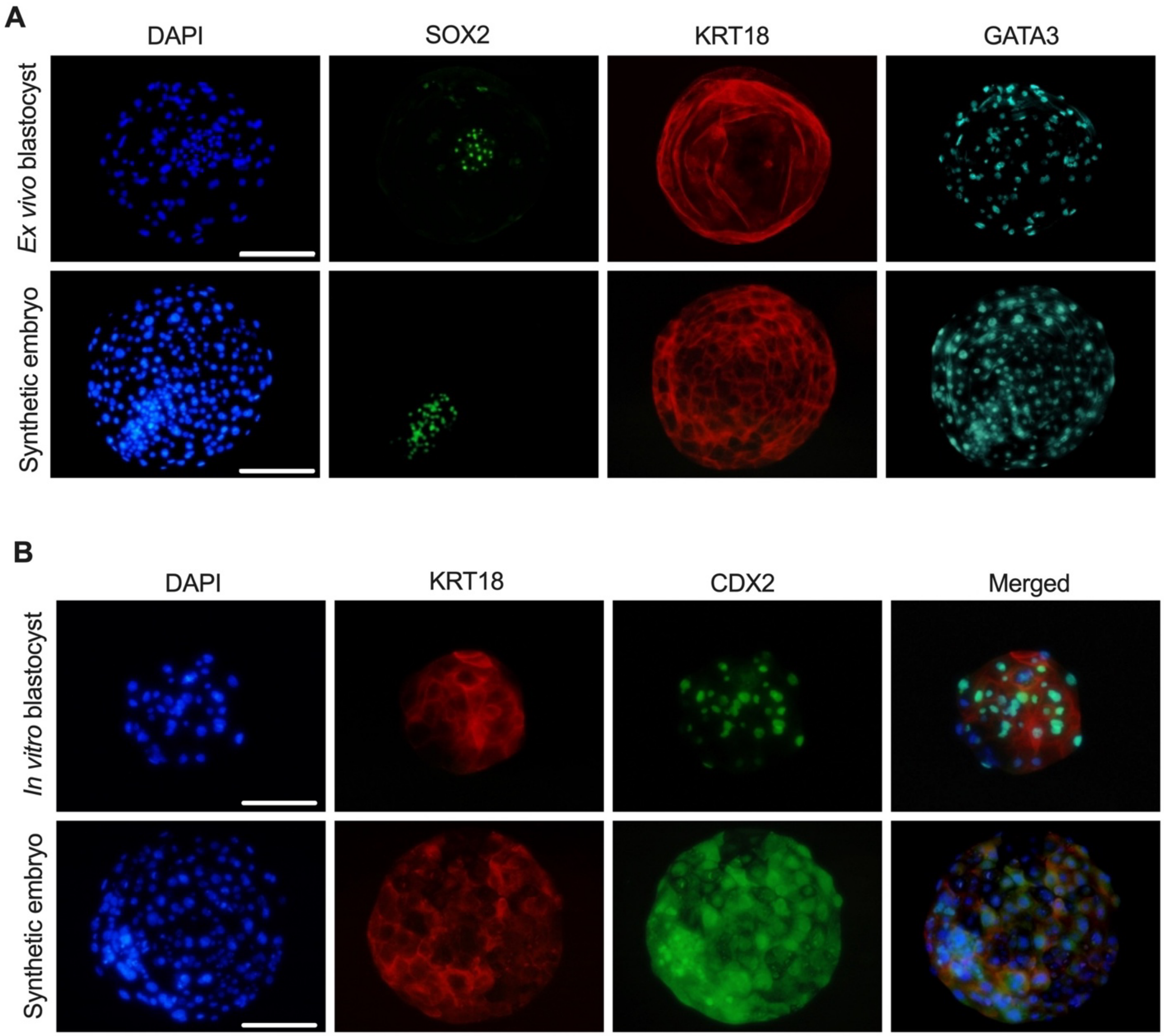


**Supplementary Figure 5. Comparative marker expression in *ex vivo* blastocysts and synthetic embryo-like structures.** Related to Figure 6

(A) Immunofluorescence analysis of *ex vivo* porcine blastocysts and synthetic embryo–like structures showing SOX2, KRT18, and GATA3 expression. Scale bars, 150 µm.

(B) Immunofluorescence comparison of *in vitro* blastocyst-like structures and synthetic embryo-like aggregates stained for DAPI, KRT18, and CDX2. Merged images illustrate the partial spatial organization of trophoblast markers in synthetic structures relative to in vitro blastocysts. Scale bars, 100 µm.

**Supplementary Tables**

**sTable 1. RT-qPCR primers**

| Gene | Forward primer (5'-3') | Reverse primer (5'-3') |
|---|---|---|
| *RN18S* | ACAAATCGCTCCACCAACTAAGA | CGGACACGGACAGGATTGAC |
| *ACTB* | GTGGACATCAGGAAGGACCTCTA | ATGATCTTGATCTTCATGGTGCT |
| *OCT3/4* | GCTGGAGCCGAACCCCGAGG | CACCTTCCCAAAGAGAACCCCCAAA |
| *SOX2* | AACAGCCCAGACCGAGTTAA | GTTGTGCATCTTGGGGTTCT |
| *NANOG* | CCCCCTTCTTCAACTCAACA | CTTCAGGCCCATAAACCTCA |
| *GATA6* | ATCACCATCACCACCCAAGT | CGCGACTCTGTAGACTGTGC |
| *SOX17* | TGGTTGAATCTTGAGGTCTGC | CAGGGTGTAGGTGTGTGATGA |
| *EOMES* | TCCCACGGATTCTCCTAGAT | AATAGCGGGCTTGAGGTAAG |
| *CDX2* | TCGCCCACAAATGTTCACCAAC | TCCAACCGCACCTGTCTTTACC |
| *GATA3* | AAAGAGAGAGAGACGGAGAGAG | CGAGGAGCAGAGAGGAGAA |
| *ETS2* | CGACAAGAACATCATCCACAAG | GATGGCGTGCAGTTCCTC |
| *ELF5* | GCTTGAAAACAAGTGGCATC | TCTTCCTTTGTCCCCACATC |
| *TEAD4* | CATGATCATCACCTGCTCCA | AAGTTCTCCAGCACGCTGTT |
| *HAND1* | ACATCGCCTACCTGATGGAC | TAACTCCAGCGCCCAGACT |
| *KRT8* | TCTGGGATGCAGAACATGAG | GGCTGTAGTTGAAGCCTGGA |
| *KRT18* | GCAAGTTCTGTGGACAATGC | GCCAGCTCCGTCTCATACTT |
| *NES* | TTCCAAGGCTTCTCTCAGCATCT | GCTCTTCAGAAAGGCTGGCATA |
| *PAX6* | CAGAGAAGACAGGCCAGCAA | GGCAGAGCACTGTAGGTGTT |
| *ZIC1* | CCAACGTGGTTAATGGGCAG | TAGTGCTCTGAACGGGGA |
| *DES* | GGCTCAGTACGAGACCATCG | GCATCGATCTCGCAGGTGTA |
| *CD44* | CAGTCACAGACCTGCCTAATAC | GGTTGGTCCTGTATTCACCTT |
| *VIM* | GTACCGGAGACAGGTGCAGT | TTCCACGGCAAAGTTCTCTT |
| *HNF4a* | CTCAGCAACGGACAGATGTG | CAGGAGCTTGTAGGGCTCAG |

**sTable 2. Antibody information**

| Antibody | Manufacturer | Species | Dilution ratio |
|---|---|---|---|
| SOX2 | Invitrogen, 14981180 | Rat | 1:100 |
| OCT4 | Stemgent, 09-0023 | Rabbit | 1:100 |
| NANOG | Abcam, ab80892 | Rabbit | 1:100 |
| SSEA4 | Invitrogen, 41-4000 | Mouse | 1:200 |
| CDX2 | Abcam, ab76541 | Rabbit | 1:100 |
| AP-2γ | Santa cruz, sc-53162 | Mouse | 1:50 |
| GATA3 | Abcam, ab182747 | Rabbit | 1:200 |
| GATA3 | R&D, AF2605 | Goat | 1:100 |
| GATA6 | R&D, AF1700 | Goat | 1:100 |
| KRT7 | Abcam, ab68459 | Rabbit | 1:200 |
| KRT7 | Santa cruz, sc-70936 | Mouse | 1:100 |
| KRT18 | Abcam, ab41825 | Mouse | 1:200 |
